# The chemical chaperone 4-phenylbutyric acid rescues molecular cell defects of *COL3A1* mutations that cause vascular Ehlers Danlos Syndrome

**DOI:** 10.1101/2024.06.20.599980

**Authors:** Ramla Omar, Michelle Lee, Laura Gonzalez-Trueba, Spyridonas Lianos, Snoopy Hazarika, Malak A Ammar, Jennifer Cassels, Alison M. Michie, Neil J Bulleid, Fransiska Malfait, Tom Van Agtmael

## Abstract

**Purpose:** Vascular Ehlers Danlos Syndrome (vEDS) is a connective tissue disorder caused by *COL3A1* mutations for which there are no treatments due to a limited understanding of underlying mechanisms. We aimed to address this critical knowledge gap, focusing on collagen folding, to establish if targeting protein folding represents a potential therapeutic approach.

**Methods:** We performed a mechanistic analysis of two novel *COL3A1* glycine mutations, G189S and G906R, using primary patient fibroblast cultures, and performed pre-clinical proof-of-concept treatments using FDA-approved chemical chaperones targeting protein folding and/or degradation.

**Results:** *COL3A1* mutations caused secretion of misfolded collagen III and intracellular collagen retention, leading to matrix defects and endoplasmic reticulum (ER) stress, with increased severity for the more C-terminal mutation. Promoting ER protein folding capacity through the chemical chaperone 4-phenylbutyric acid rescued the ER stress, thermostability of secreted collagen, matrix defects and apoptosis. Optimising treatment duration and dosage helped overcome allele-dependent treatment efficacy. In contrast, protein degradation alone or combined with targeting protein folding did not increase efficacy.

**Conclusion:** ER stress is a molecular mechanism in vEDS that can be influenced by the position of *COL3A1* mutation, and promoting protein folding is a putative mechanism-based therapeutic approach that can rescue intra- and extracellular defects.

## Introduction

Vascular Ehlers Danlos Syndrome (vEDS) (OMIM # 130050) is a rare heritable connective tissue disorder caused by heterozygous mutations in the gene *COL3A1* that encodes the alpha 1 chain of collagen III, α1(III) ^1^. VEDS is a multi-systemic disorder that significantly reduces life expectancy mostly due to dissection and rupture of arteries, intestine and gravid uterus ^2^. Other features include translucent, thin skin that tears and bruises easily and has delayed wound healing, early-onset varicose veins, small joint hypermobility, tendon- and muscle ruptures, pneumo(hemo)thorax, carotid-cavernous fistula, and characteristic facial features ^3^.

Collagen III is a major fibrillar collagen in the extracellular matrix (ECM) that is highly expressed in soft tissues with elastic properties including dermis, blood vessels, and gastro-intestinal tract ^4–6^. It is folded in the endoplasmic reticulum (ER) where three α1(III) chains interact to form a triple helical collagen III molecule in a zipper like fashion from C- to N-terminal end ^7,8^. The triple helical collagenous domain is composed of Gly-Xaa-Yaa repeats with every third amino acid being a glycine, and ∼66% of *COL3A1* mutations affect glycines ^9,10^. Mutations in other collagens reduce extracellular collagen levels and induce ER stress with activation of the unfolded protein response (UPR) that can be targeted by FDA- and EMA-approved small compounds ^11–17^. Although the first *COL3A1* mutations were identified ∼40 years ago major gaps in our understanding of their molecular mechanism remain ^1^. In particular, while ECM defects are a defining feature the impact of glycine substitutions, which account for 95% of mutations, on protein folding remains unclear.

This gap in our mechanistic knowledge is hindering the development of mechanism-based treatments. The only treatment for vEDS, in Europe, is the beta blocker celiprolol and whilst it has some efficacy ^18^, it is not well tolerated (1/3 patients do not tolerate recommended dose) ^19^ and was declined FDA approval. Experimental treatments have focused on targeting more downstream pathways or modulating known risk factors that predispose to vascular rupture such as blood pressure ^20–22^ ^1^.

Inhibiting PKC/MEK/ERK signalling (e.g. via cobimetinib), improved survival ^20^ in some but not all mouse models harbouring *Col3a1* mutations ^21^. Considering the multi-systemic nature of the disease that affects different cell types, targeting single downstream pathways may only be effective for particular cell types and/or mutations, as found for celiprolol treatment in mice ^20,22^. This underscores the need for complementary approaches that directly target upstream molecular insults of the mutations.

Here, using primary patient fibroblasts we set out to address the impact of two novel *COL3A1* glycine mutations, *COL3A1^+/G906R^* and *COL3A1^+/G189S^,* on collagen folding and establish the efficacy of FDA-approved compounds that target protein folding or degradation on the cell phenotype. Our data show that glycine *COL3A1* mutations lead to secretion of mutant collagen III while also differentially activating the UPR. Importantly, the FDA-approved chemical chaperone 4-phenylbutyric acid (PBA) rescues both of these molecular insults and improves cell viability. We also interrogated dosage and duration as treatment parameters and establish that targeting protein folding using chemical chaperones represents a potential therapeutic strategy for vEDS.

## Materials and Methods

### Cell culture and drug treatments

Primary fibroblast cultures were established from vEDS patients at the Centre for Medical Genetics, Ghent University Hospital, and controls are ethnically matched primary dermal fibroblasts (TCS Cell Works (UK) and Lonza (UK)). The research was covered by the University of Glasgow CMVLS ethics committee Ref 200200029. Cells were maintained in DMEM, 10-15% FBS and 1% penicillin/streptomycin in 37°C. For experiments cells were cultured in DMEM containing 10% FBS, 1% penicillin/streptomycin and 0.25mM ascorbic acid for 72 hours to promote collagen expression and folding ^23^. Cells were incubated with 4-PBA (PCI synthesis), tauroursodeoxycholic acid (TUDCA, Merck) or carbamazepine (CBZ, Merck) for the last 24 or 72 hours. To assess cell proliferation 30,000 cells were plated and counted using a haemocytometer at fixed intervals.

### Western Blotting

Protein extracts were prepared in RIPA buffer containing protease (Complete Mini, Roche) and phosphatase (PhosSTOP, Roche) inhibitors. Protein samples were prepared and denatured in Laemmli Buffer and SDS-PAGE was performed under reducing conditions (Mini-PROTEAN® Tetra, Bio-Rad) before transfer onto nitrocellulose membranes. Following blocking (5% milk/BSA) membranes were incubated with primary antibody overnight at 4°C (BIP [1:40,000 BD Transduction 610979], ATF4 [1:1000, Santa Cruz Biotechnology sc-200], ATF6 [1:1000 Abcam ab122897], Collagen III [1:1000, Abcam ab7778], Phospho-eIF2α (Ser51) [1:1000, Cell Signalling Technology 9721], eIF2α [1:1000, Cell Signalling Technology 9722], ubiquitin [1:1000, Santa Cruz Biotechnology P491], LC3B [1:500, Novus Biologicals 1251A], Tubulin [1:40,000 Sigma T5168]). Following HRP-conjugated secondary antibody incubation (1:1000, Cell Signalling Technology), membranes were incubated Luminata Forte Western HRP substrate (Millipore) before visualization using a BioRad ChemicDoc XRS+ or X-OMAT-Film processor using Hyperfilm™ ECL (GE Healthcare).

### Thermal stability of collagen

30 µl conditioned medium from cells was treated with 1 μl 0.05 % trypsin at 46°C for 30sec, 1min or 3min. Trypsin was inactivated by adding 1 μl 0.1% trypsin inhibitor (Sigma) and denaturation at 95°C for 5min. The samples were then analyzed using western blot.

### qRT-PCR

Cells were incubated with Trizol (ThermoFisher) and RNA was extracted and resuspended in nuclease free water as per manufacturer’s Instruction. RNA samples were treated with DNA-free™ DNA Removal Kit (ThermoFisher) as per manufacturer’s Instruction, and subsequently spiked with luciferase (50pg/µg RNA). Following cDNA synthesis (High Capacity cDNA Reverse Transcription Kit (Thermofisher), qRT-PCR was performed using Power Up SYBR green Master mix (Invitrogen; 7900HT Fast Real-Time PCR Applied Biosystems). Samples were normalized to 18S RNA and/or luciferase and relative mRNA levels were calculated using the 2^-ΔΔ*CT*^ method. Primer sequences: CHOP (GCGCATGAAGGAGAAAGAAC, TCTGGGAAAGGTGGGTAGTG), IRE1 (CGGGAGAACATCACTGTCCC, CCCGGTAGTGGTGCTTCTTA), COL3A1 (TGGTCTGCAAGGAATGCCTGGA, TCTTTCCCTGGGACACCATCAG), 18S (AGTCCCTGCCCTTTGTACACA, CGATCCGAGGGCCTCACTA), luciferase (GCTGGGCGTTAATCAGAGAG, GTGTTCGTGTTCGTCCCAGT).

### Analysis of apoptosis

Cultured cells were collected and apoptosis measured using the FITC Annexin V Apoptosis Detection Kit I (BD biosciences). 1x 10^5^ cells were stained and cells (6000 events/sample) were analysed in triplicate using a BD FACS Canto II and FlowJo Software. Unstained, FITC stained only, PI stained only and positive control (DMSO-induced apoptosis) cells were used to calibrate the machine.

**Immunocytochemistry** was performed as described ^16^. Cells were fixed (acetone, 10% methanol, or 4% PFA for 20 min at −20 °C), washed in PBS, and incubated with 0.05 M KCL/0.05 M HCL for 15 min at RT for antigen retrieval. Following blocking, cells were incubated with primary antibodies in 10% goat serum 4°C overnight (collagen III [1:300, Abcam ab778], PDI [P4HB 1:200, Abcam ab2792]. Cells were washed in PBS, incubated with secondary antibody (Cy2, Cy3 conjugated, 1:500 Jackson Immunoresearch) in 1% goat serum. Following washes, cells were incubated with DAPI, washed, and mounted (Fluoromount-G, Invitrogen) prior to imaging on a Nikon eclipse Tg2 fluorescence microscope. For extracellular collagen III staining 50,000 cells were seeded on coverslips, cultured for 24 hours prior to 72 hour culturing in media containing 0.25 mM ascorbic acid with/without PBA. Coverslips were washed with PBS prior to decellularization by incubating with 20 mM NH_4_OH for 30 minutes at RT, followed by washes with dH_2_O then PBS. The matrix was fixed with 4% PFA (10 minutes, RT), samples were washed with PBS, blocked in 10% goat serum in PBS (1 hour RT), and washed thrice in PBS. Samples were incubated with anti-collagen III antibody (1:100, Abcam ab7778) overnight at 4°C. Following washes, samples were incubated with secondary antibody (1:300, goat-anti-rabbit Alexa Fluor 488, ab150077, RT, 1 hour), mounted on slides and imaged using a Nikon Eclipse Ts2 microscope with Nikon DS-Fi3 camera using NIS Elements Software (Nikon). Matrix deposition was quantified by measuring integrated density (Image J). Values were averaged across 5 pictures taken per coverslip per n number.

**Statistical Analysis** was performed using GraphPad Prism software. Unpaired Student t-test, One-Way ANOVA with post hoc correction, or 2-way ANOVA were used as appropriate. p-value <0.05 was considered statistically significant.

## Results

### *COL3A1* glycine mutations have a quantitative and qualitative effect in vEDS

To explore pathomolecular mechanisms of *COL3A1* glycine mutations, we established primary dermal fibroblasts cultures of two vEDS patients. Both mutations affect glycine residues of the Gly-Xaa-Yaa repeat in α1(III) with the more N-terminal mutation in exon 6 leading to a glycine to serine substitution (G189S), and the more C-terminal mutation altering glycine to arginine in exon 31 (G906R) (Fig. 1A-B). The G189S mutation is absent in gnomAD and ClinVar, but G189R (rs587779507) has been reported on dbSNP and ClinVar. Analysis using Variant Effect Predictor software in Ensembl (SIFT, PolyPhen, CADD score) classified both variants as deleterious (Table 1), supporting the causality of these two novel mutations.

**Figure 1.**
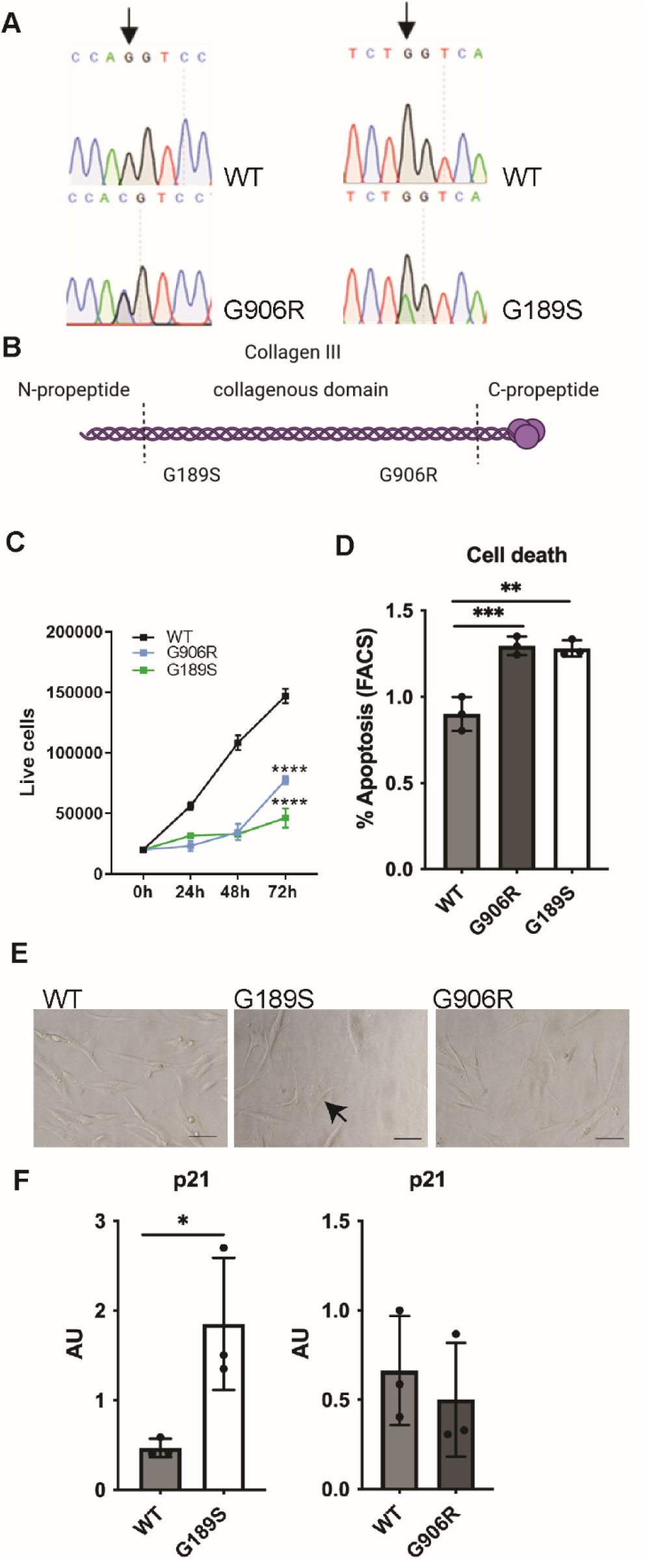
Identification and functional defects of *COL3A1* mutations. (A) Sanger sequence traces of PCR products from DNA of primary patient fibroblasts cultures, covering COL3A1 G906R and G189S mutation. WT: control primary dermal fibroblast. Black arrow indicates position of heterozygous mutation. (B) Diagram showing position of mutation within the collagen III protein and protein domain structure. (C) Cell proliferation analysis of wild type (WT), *COL3A1^G189S/+^* (G189S) and *COL3A1^G906/+^* (G906R) cells (n= 3, Two-Way ANOVA). (D) Percentage apoptotic cells determined by FACS (Annexin V, propidium iodide positive, FACS scatter plot provided in Supplemental Figure 1)) of wild type (WT), *COL3A1^G189S/+^*(G189S) and *COL3A1^G906/+^*(G906R) cells (n= 3; One-way ANOVA with Dunnett’s multiple comparison test). (E) Brightfield microscopy shows flattened irregular shaped enlarged cells particularly in *COL3A1^G189S/+^.* (F) qRT-PCR reveals increased mRNA levels of p21, marker of senescence, in *COL3A1^G189S/+^* cells (n=3, unpaired t-test). * p<0.05; ** p<0.01; *** p<0.001 **** p<0.0001.

**Table 1.** In silico analysis of *COL3A1* mutations. . CADD >20 and >30 indicate top 1% and top 0.1% of single nucleotide variants.

| SYMBOL | EXON | cDNA | Protein | SIFT | PolyPhen | CADD_PHRED |
| --- | --- | --- | --- | --- | --- | --- |
| COL3A1 | 6/51 | 682 | G189S | deleterious(0.01) | probably_damaging(1) | 32 |
| COL3A1 | 39/51 | 2833 | G906R | deleterious(0) | probably_damaging(0.999) | 31 |

Both mutations caused a reduction in cell proliferation (Fig. 1C) with increased apoptosis (Fig. 1D, Supplemental Figure 1) and we also observed altered morphology of mutant cells which was particularly pronounced in *COL3A1^+/G189S^* cells, looking flatter with an irregular larger “pancake” type morphology compared with wild type (WT) fibroblasts (Fig. 1E), a key morphological feature of senescent cells ^24^. These features coupled with increased levels of the senescence marker p21 in *COL3A1^+/G189S^*(Fig. 1F) ^25^, suggest senescence induction due to *COL3A1* mutations.

Non-mutually exclusive impacts of collagen mutations include secretion of mutant protein, reduced extracellular protein levels and/or protein misfolding leading to ER stress ^1,26^. To shed light on these, we determined if these mutations affect collagen III secretion. This revealed increased intracellular collagen III levels in both mutant cells, and reduced (G189S) or similar (G906R) extracellular levels (Fig 2A-B), indicating a shift towards intracellular retention. Immunostaining confirmed collagen III was retained in the ER and staining against the ER marker PDI also revealed ER enlargement (Fig. 2C), a sign of ER stress. While *COL3A1* nonsense mutations show reduced protein levels can be pathogenic, the increased severity of glycine substitutions supports a dominant negative effect ^3,9^, potentially by secreting mutant protein. We used sensitivity to trypsin digestion as a proxy of the folding quality of triple helical secreted collagen III. This revealed that secreted collagen III from mutant cells is digested more rapidly (Fig. 2D) and that both glycine mutations enable secretion of misfolded collagen III. This is associated with an altered more punctate appearance of the deposited collagen III network with apparently less fibrils (Fig. 2E). These data show that collagen III mutations act via quantitative and qualitative effects by reducing levels of extracellular collagen III coupled with secreting mutant less stable collagen III.

**Figure 2.**
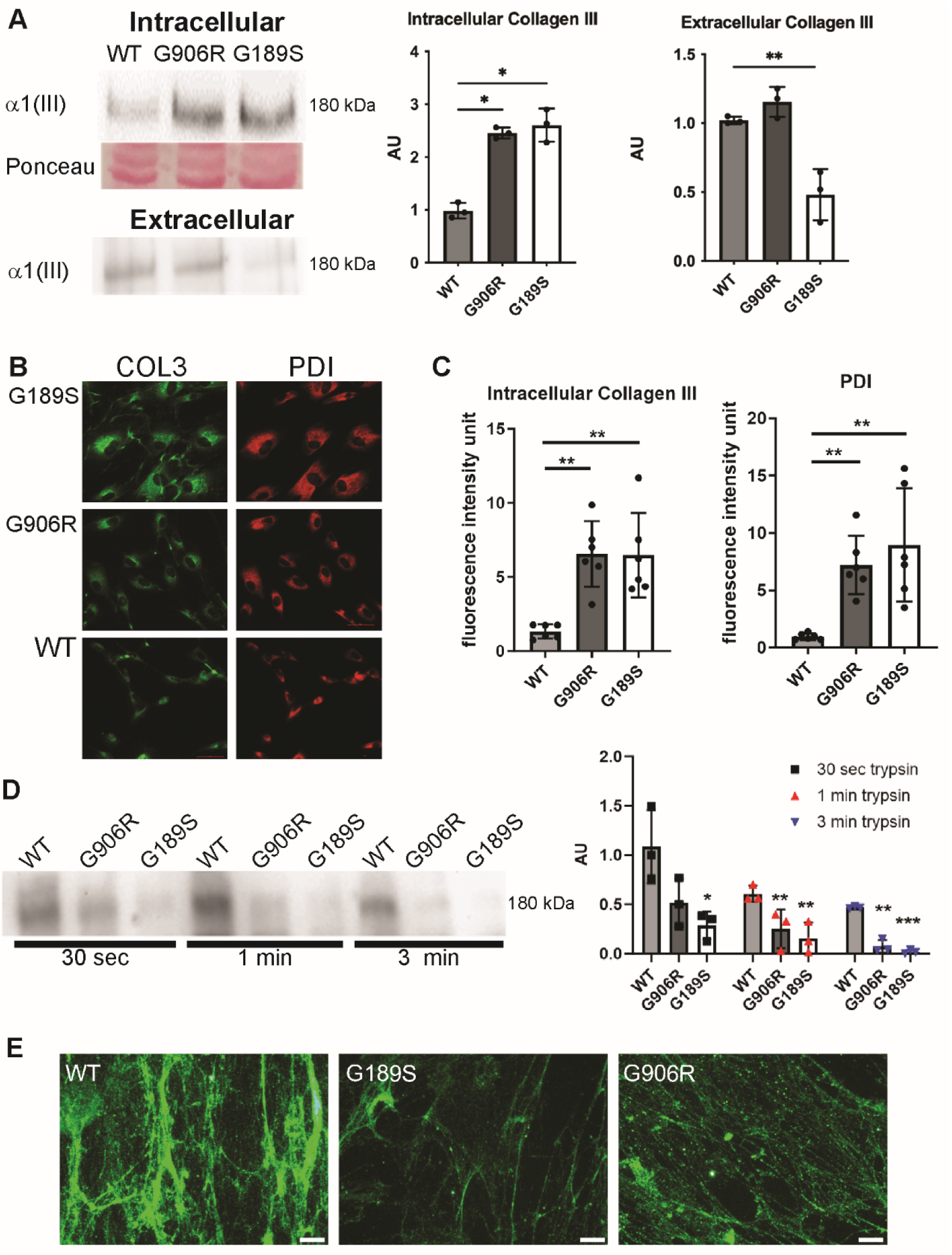
COL3A1 mutations affect collagen III protein handling. (A) Western blotting against collagen III on cellular protein lysate (intracellular) and conditioned media (extracellular) of wild type (WT), *COL3A1^G189S/+^*(G189S) and *COL3A1^G906/+^*(G906R) cells reveals intracellular retention. Ponceau total protein stain used as protein loading control. Quantification provided on right hand side. (n=3). (B) Immunostaining against collagen III (green) and PDI (red, ER marker) on wild type (WT), *COL3A1^G189S/+^*(G189S) and *COL3A1^G906/+^* (G906R) cells showing collagen III retention in ER in mutant cells. Size bar 50µm (C) Image J analysis using integrated density of fluorescence staining reveals ER retention and enlarged ER area (n=6). (D) Western blotting against collagen III on conditioned media that has been subjected to trypsin digestion (proxy of collagen triple helix folding) reveals secretion of mutant misfolded protein. Duration of trypsin digest is indicated. Quantification of western blot on right-hand side (n=3). (E) Immunostaining against collagen III (green) of decellularized matrix shows less developed collagen III ECM network and punctate appearance in mutant cells (n=3). A,C,D One Way ANOVA with Dunnett’s multiple comparison test * p<0.05; ** p<0.01; *** p<0.001 .

### vEDS mutations activate the unfolded protein response

Misfolding of secreted proteins can lead to ER stress and activation of the UPR, which consists of three signalling arms mediated by PERK, IRE1 and ATF6 ^27^. IRE1 activation leads to “splicing” of the mRNA XBP1, while PERK phosphorylates eIF2α and causes upregulation in mRNA translation of ATF4 ^27^. *COL3A1* mutant cells had increased levels of the ER chaperone BIP (Fig. 3A-B) with a more prominent activation of the IRE1 arm, in particular in *COL3A1^+/G906R^* (Fig. 3C). Similarly, while both mutations caused EIF2α phosphorylation, only *COL3A1^+/G906R^* showed increased levels of ATF4, although a trend was observed in *COL3A1^+/G189S^*cells (Fig. 3A-B). We did not detect activation of the ATF6 arm (Fig. 3A-B), indicating *COL3A1* mutations do not activate all arms of the classical ER stress response. The increased levels of CHOP (Fig. 3D) in *COL3A1^+/G906R^* cells but not *COL3A1^+/G189S^* support presence of ER stress-associated apoptosis. ER stress can activate protein degradation pathways to decrease misfolded protein levels in the ER ^27^. Western blotting revealed activation of proteasomal degradation pathways, shown by increased levels of poly-ubiquitinated proteins, but not of autophagy, probed by assessing LC3I to LC3II conversion (Fig 3E).

**Figure 3.**
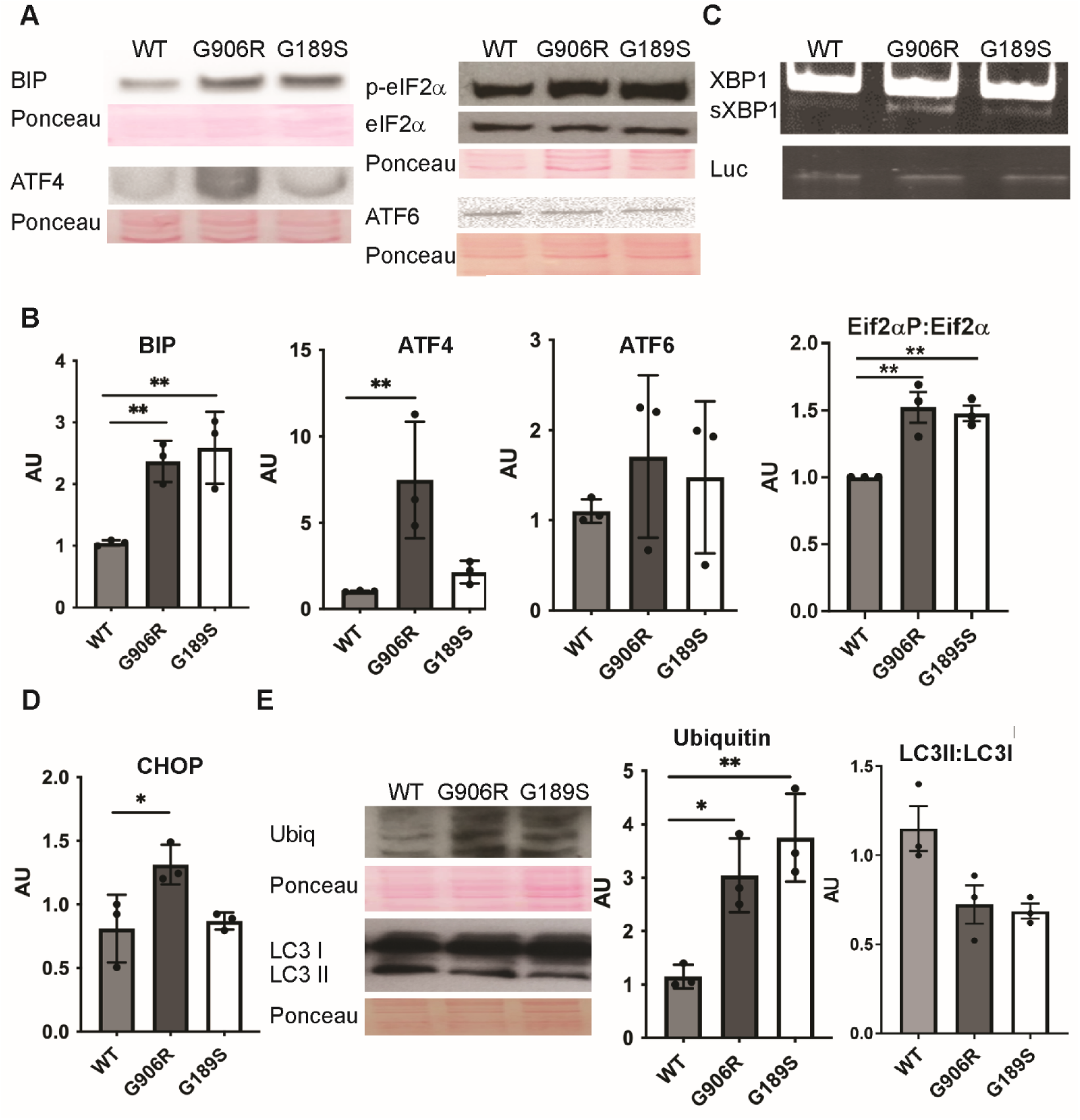
*COL3A1* mutations activate ER stress. (A) Western blotting against ER stress markers in wild type (WT), *COL3A1^G189S/+^* (G189S) and *COL3A1^G906/+^* (G906R) cells. eIF2α: total eIF2α; p-EIF2α: phosphorylated EIF2α. Ponceau staining loading control. (B) Densitometry analysis of bands in (A) shows UPR activation in mutant cells. (C) Representative gel of RT-PCR showing splicing of XBP1 by IRE1. Spliced XBP1 (sXBP1), unspliced XBP1 (XBP1), Spiked luciferase (Luc) used as loading control (see Materials and Methods for further details). (D) Measurement of CHOP mRNA levels by qRT-PCR (n=3). (E) Analysis of proteasome and autophagy levels by western blotting against ubiquitinated proteins and LC3BI-II. Densitometry analysis by Image J provided in graphs on right-hand side (n=3). B, D, E One Way ANOVA with Dunnett’s multiple comparison * p<0.05; ** p<0.01.

These data establish that *COL3A1* mutations induce differential UPR activation with a more extensive and chronic ER stress due to the C-terminal *COL3A1^+/G906R^*mutation.

### Targeting protein folding using PBA rescues intracellular phenotypes

Conceptually, pharmacologically targeting collagen folding and/or mutant protein degradation could modulate both the ER stress and ECM defects by promoting collagen secretion ^15,16,28^ and/or secretion of better-folded collagen. This could represent an avenue for rescuing both extra- and intracellular effects and be effective across different *COL3A1* mutations and tissues. This would overcome the genotype-dependent efficacy of recently proposed strategies that target more downstream mechanisms or blood pressure ^20–22^. The availability of FDA-approved compounds, including PBA, TUDCA and CBZ, that target protein folding or degradation is particularly interesting as these are well-tolerated with good safety records ^29^, and repurposing FDA/EMA-approved compounds is an attractive cost-effective strategy for developing treatments for rare diseases ^30^.

We therefore set out to investigate their efficacy on *COL3A1* mutations by first incubating cells for 24 hours with different concentrations of PBA (1mM, 5mM, 10mM), TUDCA (10μM, 100μM, 1mM) and CBZ (10μM, 20μM, 1mM) to assess the highest concentration that is tolerated by control primary dermal fibroblasts. Analysis revealed reduced viability with 10mM PBA and 1mM CBZ, while cell survival was not impacted by TUDCA treatment up to 1mM (Supplemental Figure 2).

As PBA can alleviate cellular defects due to mutations in other collagen types ^15–17,31^, we first investigated its efficacy on vEDS fibroblasts. Incubating cells in 5mM PBA for 24 hours reduced BIP protein levels in *COL3A1*^+/G189S^ but not *COL3A1^+/G906R^* cells (Fig. 4A-B). To establish if the reduction in BIP levels is shared with other chemical chaperones we incubated vEDS cells with 500µM TUDCA. This also reduced levels of BIP in *COL3A1*^+/G189S^ cells (Supplemental Figure 3), supporting that the reduction in BIP levels is at least in part due to effects on protein folding. In contrast, CBZ did not alter BIP protein levels or the phospho-Eif2α:total Eif2α ratio, revealing no ER stress reduction (Supplemental Figure 4) through promotion of protein degradation, in contrast to what is achieved in *COL10A1* mutations ^14^. These data show that chemical chaperones can rescue ER stress caused by *COL3A1* mutations but with allelic-specific effects as the C-terminal mutation was more resistant to treatment.

**Figure 4.**
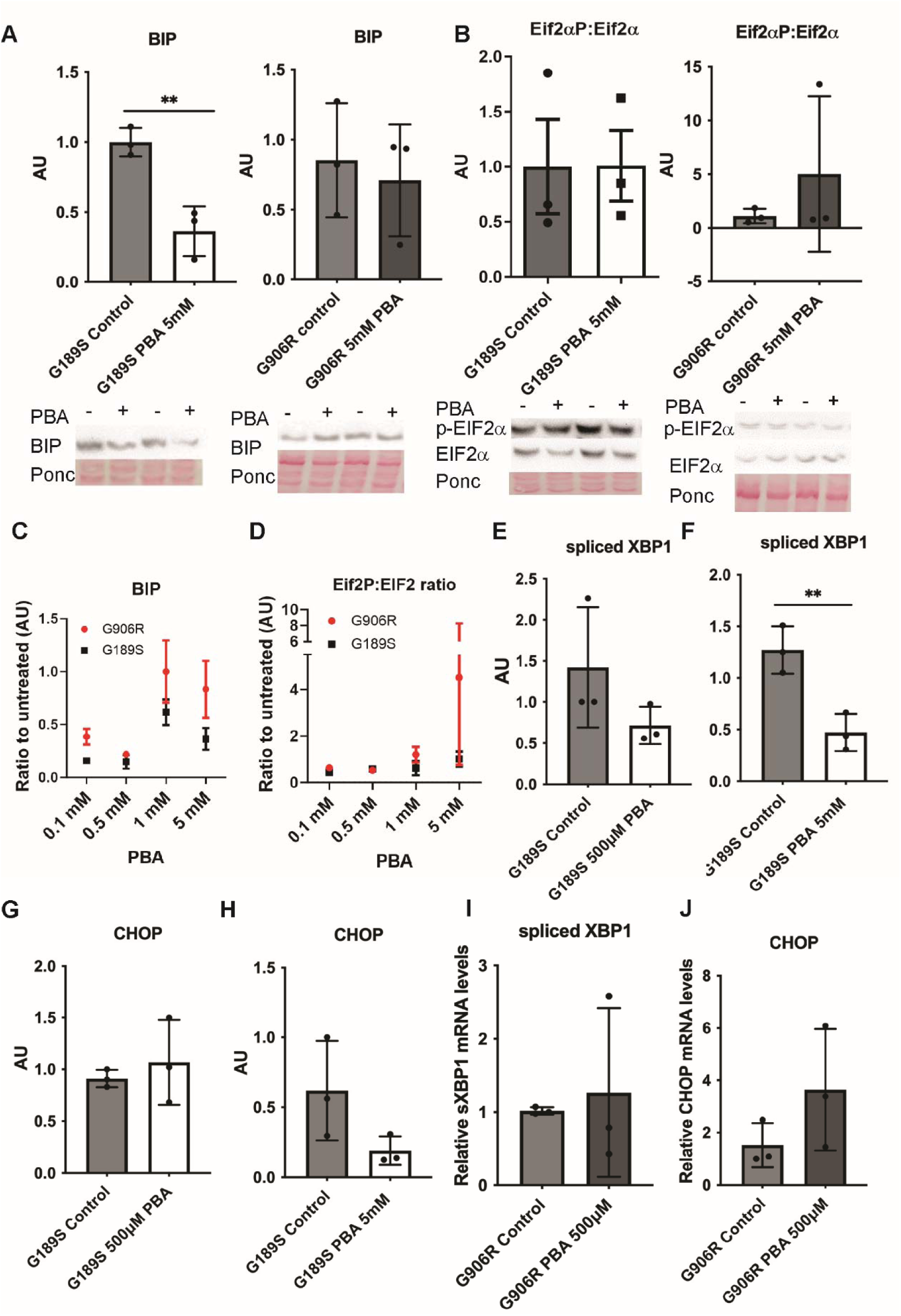
Efficacy of PBA in reducing intracellular defects of *COL3A1* mutations. (A) Western blotting against BIP in PBA-treated (PBA+ in gel) and untreated (control in graph, PBA – on gels) *COL3A1^G189S/+^* (G189S) and *COL3A1^G906/+^* (G906R) cells shows allele specific efficacy. Gels showing two biological replicates are provided below graphs of densitometry analysis using Image J (n=3, unpaired t-test). Ponc: ponceau staining as protein loading control. (B) Western blotting against total (EIF2α) and phospho-eIF2α (p-EIF2α) and graphs of densitometry analysis using Image J. (C) Graph of densitometry analysis of BIP protein levels determined by western blot (gels provided in Supplemental Figure 5) in cells incubated for 24 hours with increasing PBA concentrations (0.1 mM, 0.5 mM, 1 mM, 5mM). (D) Graph of densitometry analysis of ratio of phospho-EIF2α versus total EIF2α protein levels determined by western blot (gels provided in Supplemental Figure 5) in cells incubated for 24 hours with different PBA concentrations. (E-F) qRT-PCR analysis of spliced XBP1 in untreated *COL3A1^G189S^*^/+^ cells (control) and cells treated with 500 µM and 5mM PBA for 24 hours. (n=3). (G-H) qRT-PCR analysis of CHOP in untreated *COL3A1^G189S^*^/+^ cells (control) and cells treated with 500 µM and 5mM PBA for 24 hours (n=3). (I-J) qRT-PCR analysis of spliced XBP1 and CHOP in untreated *COL3A1^G906R^*^/+^ cells (control) and treated with 500 µM for 24 hours. (n=3). A-B, E-J unpaired t-test * p<0.05; ** p<0.01.

### PBA dosage impacts treatment efficacy

Treatment dosage is an important consideration for any future treatments and given the allele specific effects, we set out to explore the impact of PBA dosage on its efficacy. Cells were incubated with four PBA concentrations for 24 hours and ER stress was measured. This revealed PBA reduces levels of BIP and the PERK pathway in *COL3A1^+/G189S^* across the different concentrations (Fig 4C-D, Supplemental Figure 5). In contrast only lower PBA concentrations reduced ER stress marker levels in *COL3A1^+/G906R^* (Fig. 4C-D, Supplemental Figure 5).

We further characterised the effects of 24 hour incubation with 5mM and 500µM PBA on *COL3A1^+/G189S^* which revealed that 5 mM PBA significantly reduced spliced XBP1 levels with a trend towards lower CHOP mRNA levels in COL3A1^+/G189S^ (Fig. 4E-H). This provides evidence for potential higher efficacy of 5mM PBA compared to 500µM. In contrast, 500µM PBA, the concentration that showed most promising effect on BIP and EIF2α phosphorylation in *COL3A1^+/G906R^*, had no effect on CHOP or spliced XBP1 levels (Fig. 4I-J). Given the absence of impact of higher dosage on BIP, we employed 500µM PBA in *COL3A1^+/G906R^* cells.

To investigate if the lower ER stress levels were associated with reduced ER retention of collagen III, we performed western blotting on cell lysates. This supported reduced intracellular α1(III) retention in *COL3A1^+/G189S^* but not *COL3A1^+/G906R^* despite the modulated BIP levels and p-EIF2α/EIF2α ratio (Fig. 5A-B). Given the limited efficacy of 500µM PBA in *COL3A1^+/G906R^*, we next explored the impact of increased treatment duration for *COL3A1^+/G906R^*. This revealed that a 72 hour incubation reduced intracellular collagen III levels (Fig. 5A-B), which was confirmed by immunostaining showing reduced ER retention of collagen III and ER area (Fig. 5C-D). These data establish increased efficacy with a longer PBA incubation. To explore why 72-hour and not 24-hour incubation with PBA reduced collagen III levels in *COL3A1*^+/G906R^, we determined *COL3A1* mRNA levels as PBA can also have HDAC inhibitor activity ^29^. This revealed genotype dependent effects with increased mRNA levels with 24 hour 500μM PBA but not 72-hours for *COL3A1*^+/G906R^, and reduced *COL3A1* mRNA levels in *COL3A1*^+/G189S^ (Supplemental Figure 3). This apparent pulse in expression could explain the lack of intracellular collagen III reduction with 24 hour PBA incubation in *COL3A1*^+/G906R^ cells. We also explored if a combinatorial treatment that simultaneously targeted protein folding and degradation increased efficacy for *COL3A1*^+/G906R^. Coupling 500µM PBA with 20µM CBZ for 24 hours did not reduce intracellular collagen III levels (Supplemental Figure 6A). Combined, these data support that PBA can rescue the ER stress due to *COL3A1* missense mutations and that dosage and treatment duration are important treatment parameters to help overcome allele dependent intracellular effects.

**Figure 5:**
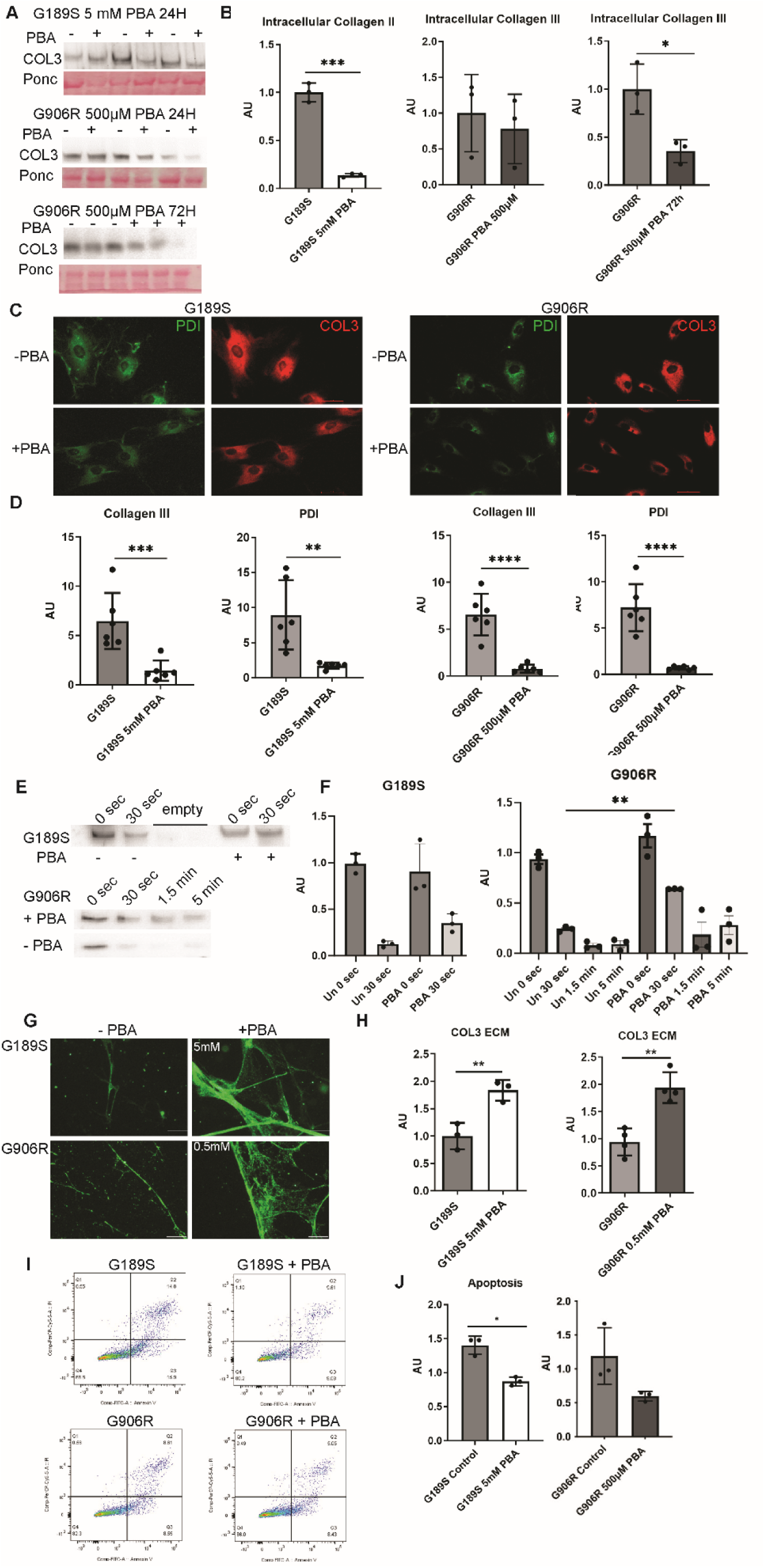
PBA dosage and rescue of cell and ECM defects. . (A) Western blotting against intracellular collagen III on untreated (-) and PBA-treated cells (+). *COL3A1^G906R^*^/+^ cells were treated with 500 µM PBA for 24 and 72 hours. (B) Densitometry analysis of gels in (A) (n=3). (C) Immunostaining against collagen III (red) and PDI (green, ER marker) on untreated (-PBA) and PBA treated (+PBA) *COL3A1^G189S/+^* (G189S) and *COL3A1^G906/+^* (G906R) cells. *COL3A1^G189S/+^* (G189S) was treated with 5 mM PBA for 24 hours, *COL3A1^G906/+^* with 500 M PBA for 72 hours. Scale bar 50 µm. (D) Image J analysis using integrated density of fluorescence staining reveals reduced ER retention of collagen III and ER area in PBA treated cells (n=6). (E) Western blotting against collagen III on conditioned media from untreated (PBA -) and PBA (PBA +) treated cells that has been subjected to trypsin digestion. *COL3A1^G189S/+^* (G189S) was treated with 5 mM PBA for 24 hours, *COL3A1^G906/+^* with 500 M PBA for 72 hours. Duration of trypsin digest is indicated. (F) Quantification of western blot on right-hand side supports PBA increased resistance to trypsin digest (n=3). (G) Immunostaining against collagen III (green) on decellularized deposited ECM from untreated and PBA treated cells. *COL3A1^G189S/+^* (G189S) and *COL3A1^G906/+^* (G906R) cells were incubated three days with 5 mM and 500 µM PBA respectively. Scale bar 10 µm. (I) FACS scatter plot of untreated (G189S, G906R) and treated cells (+ PBA) stained with propidium iodide (PI) and Annexin V. *COL3A1^G189S/+^* (G189S) was treated with 5 mM PBA for 24 hours, *COL3A1^G906/+^* with 500 M PBA for 72 hours. (J) Graphs of apoptosis levels as determined by FACS in (I) showed PBA reduced apoptosis.

### PBA rescues extracellular defects

We next set out to determine the impact of PBA on secreted collagen III given the established role of matrix defects in vEDS ^1^. Western blotting of conditioned media indicated that PBA did not significantly increase the levels of collagen III secreted over a 24 hour period (Supplemental Figure 6B). However, the secreted collagen III was more resistant to trypsin digestion (Fig. 5F), and there was increased collagen III incorporation into the deposited ECM that showed a better-formed network (Fig. 5G). Furthermore, PBA treatment also reduced the apoptosis of patient fibroblasts (Fig. 5H). Therefore, PBA increased the quality of the secreted collagen and rescued both intracellular and extracellular sequelae as well as apoptosis due to *COL3A1* mutations.

## Discussion

Here, we provide novel insight into the molecular basis of genotype-phenotype correlation and mechanisms of vEDS by uncovering that glycine *COL3A1* mutations lead to secretion of mutant protein coupled with retention of collagen III in the ER that causes allele-specific UPR activation, apoptosis and ECM defects. Targeting protein folding using the FDA-approved chemical chaperone PBA rescues the ECM, molecular and cellular defects of these mutations and treatment duration is an important parameter to help overcome allele-specific effects of mutations. Combined these data support PBA represents a putative mechanism-based therapeutic approach for vEDS.

Glycine mutations account for the majority of *COL3A1* mutations in vEDS ^9,10^ but their molecular mechanisms and that of vEDS remain poorly understood. Genetics data and outcomes in pre-clinical treatments ^9,20,21^ support allele-specific effects but the molecular basis of any genotype-phenotype correlation remains incomplete. Our data establish that combined with secreting misfolded protein, *COL3A1* mutations induce differential UPR activation with a more extensive and chronic ER stress due to the C-terminal *COL3A1^+/G906R^*mutation that include activation of the PERK and IRE1 signalling arms, which was more resistant to treatment. This differential UPR activation may thus contribute to the basis of the genotype-phenotype correlation in vEDS. The increased severity of the *COL3A1^+/G906R^* mutation could relate to the characteristics of the mutation as replacement of glycine with a larger amino acid is associated with more severe vEDS ^9^, and arginine in G906R is larger than serine in the more N-terminal *COL3A1 G189S* mutation. Moreover, it can reflect a positional effect as more C-terminal mutations have been associated with increased disease severity in fibrillar collagen disorders ^32^. It is tempting to suggest this is due to more detrimental impact on protein folding with mutations closer to the initiation site of triple helix formation, which proceeds in a C- to N-terminal direction end ^1,7,8^. It is therefore tempting to suggest the degree of UPR activation and impact on proteostasis contributes to the basis of the mutation and disease severity.

Our UPR data provide direct evidence that ER stress and UPR activation are a feature of vEDS. This is supported by signs of ER stress in the vasculature of a vEDS mouse model ^21^, defects in ER homeostasis on transcriptomic analysis of fibroblasts ^33,34^, and a delay in protein folding due to *COL3A1* glycine mutations in a bacterial expression system ^35^. The detection of ER stress in fibroblasts but limited evidence from smooth muscle cells ^20,21^ could reflect a cell type-dependent mechanism whereby UPR activation may occur more readily due to the higher expression of ECM proteins in fibroblasts compared to smooth muscle cells (Human Protein Atlas) ^36^; Dataref: Karlsson et al., 2021).

Recent data from mouse models explored the efficacy of modulating more downstream ERK signalling mechanisms ^20^. We set out a complementary approach focusing on more upstream mechanisms that may be applicable to multiple mutations and tissues to help overcome recently observed allele specific outcomes of treatments targeting these further downstream mechanisms ^20–22^. Excitingly PBA rescued both the ER stress and improved the quality of the secreted collagen without increasing levels of secretion per se, as determined by the susceptibility to trypsin digest. This raises the intriguing prospect that PBA could rescue both cell and ECM defects. While it is necessary to extend these data into vascular cells and mouse models, the ability to ameliorate upstream molecular pathological sequelae is particularly appealing for genetic disorders, such as collagenopathies and vEDS, where most patients have non re-occurring mutations ^9,37,38^. Moreover, collagens such as collagen III are widely expressed ^1^ and different cell types likely have distinct downstream responses to the initial underlying pathomolecular event. Thus, the ability to modulate these initiating pathomolecular mechanisms is an attractive approach for future therapies.

Our data also showed that promoting protein degradation by CBZ was not able to reduce the ER stress, in contrast to *COL10A1* mutation in chondrodysplasia type Schmidt where CBZ is currently being used in a clinical trial ^14^. A combination of PBA and CBZ was also not effective. Combined with data from collagen I and IV ^15,31,39^, this supports that targeting protein folding rather than degradation has more efficacy across different collagen types. This may help further stratification of collagenopathies into arms for any future mechanism-based treatments and trials targeting shared mechanisms.

In conclusion, these data establish that *COL3A1* mutations in vEDS cause ER stress via IRE1 and PERK activation, and also enable secretion of mutant protein, increasing our mechanistic insight into vEDS. Moreover, these defects were rescued by the FDA-approved chemical PBA, indicating this represents a putative therapeutic strategy that can overcome allele-specific disease mechanisms.

## Acknowledgements

We would like to thank the patients and their families for their participation in this project. This work was funded by a University of Glasgow MVLS DTP scholarship to RO; a BHF studentship to LGT (FS/4yPhD/F/20/34127); a MRC Project grant (MR/R005567/1), Stroke Association Programme Grant (16VAD_04) and Heart Research UK Translational Award (RG 2664/17/20) to TVA. FM is a senior clinical investigator of the Research Foundation Flanders (18423181N), and this work funded by Ghent University (GOA019-21).

## Author Contributions

Study design and concept: RO, NB, FM & TVA; Experimental work RO, ML, LGT, SH, MAA, TVA, SL, JC (supervised by AMM); Data Analysis: RO, ML, LGT, FM, TVA; Writing of the manuscript: all authors.

