## Supplemental Data for "The chemical chaperone 4-phenylbutyric acid rescues molecular cell defects of *COL3A1* mutations that cause vascular Ehlers Danlos Syndrome"

**Supplemental Figures**

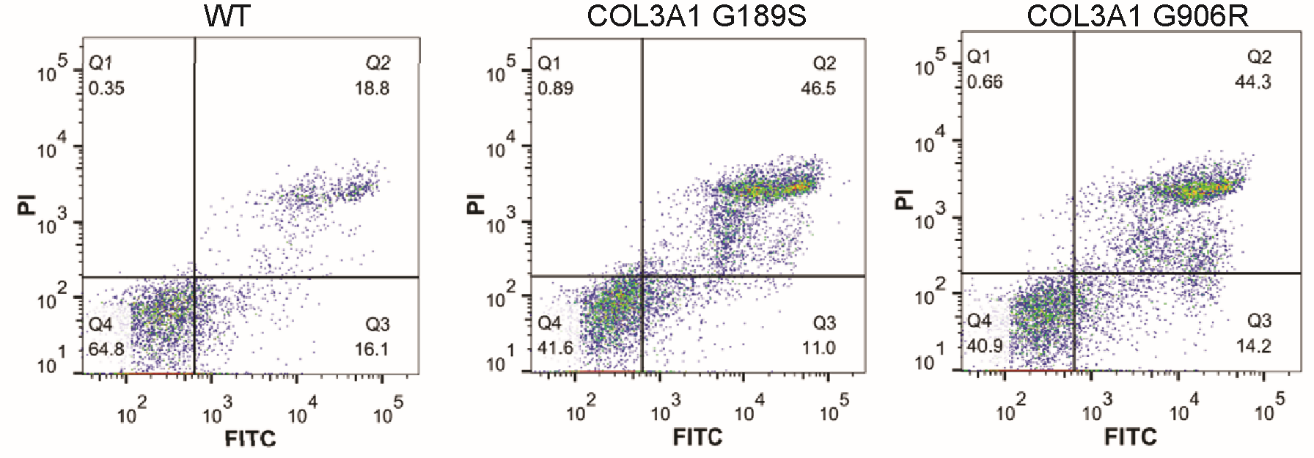

***Supplemental Figure 1. COL3A1* mutations induce apoptosis**. Scatter plot of FACS data shown in Figure 1 on wild type (WT) cells and *COL3A1^G189S/+^* (COL3A1 G189S) and *COL3A1^G906R/+^* (COL3A1 G906R) cells stained with propidium iodide (PI) and annexin V (FITC).

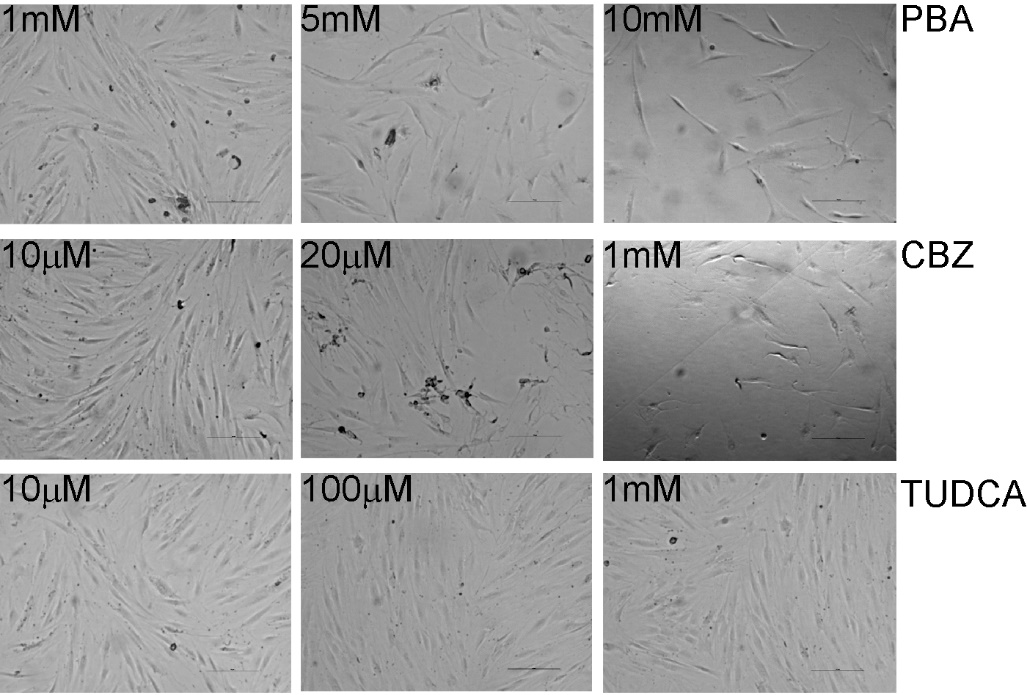

**Supplemental Figure 2. Impact of treatments on cell viability. Phase contrast** Microscopy images of wild type primary dermal fibroblasts incubated for 24 hours with varying concentrations of PBA, TUDCA and CBZ. Scale bar 100 µm

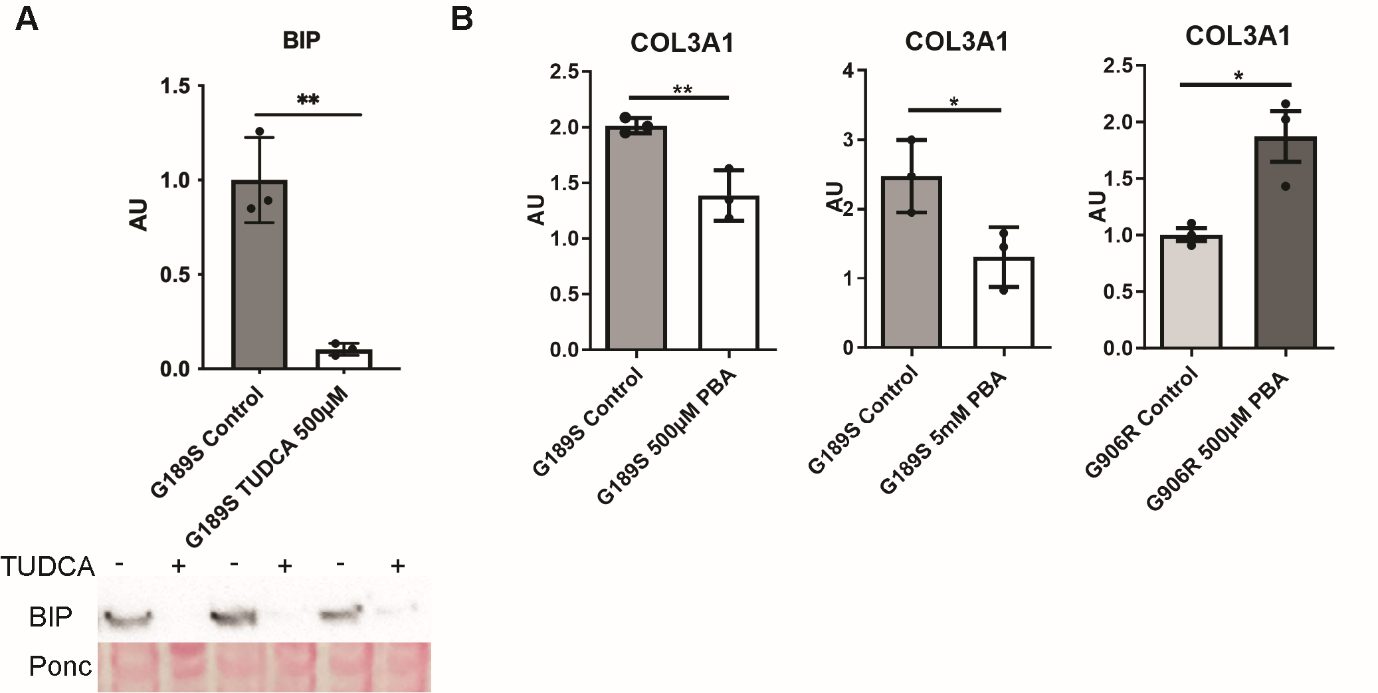

**Supplemental Figure 3 Efficacy of TUDCA on ER stress and PBA on COL3A1 mRNA levels** (A) *COL3A1^G189S/+^* (G189S) cells and cells treated with TUDCA for 24 hours, shows reduced levels of ER stress marker BIP. N=3, unpaired t-test ** p<0.01 (B) *COL3A1^G189S/+^* (G189S) and *COL3A1^G906R/+^* (G906R) cells, and cells treated with 0.5 mM or 5mM PBA for 24 hours.

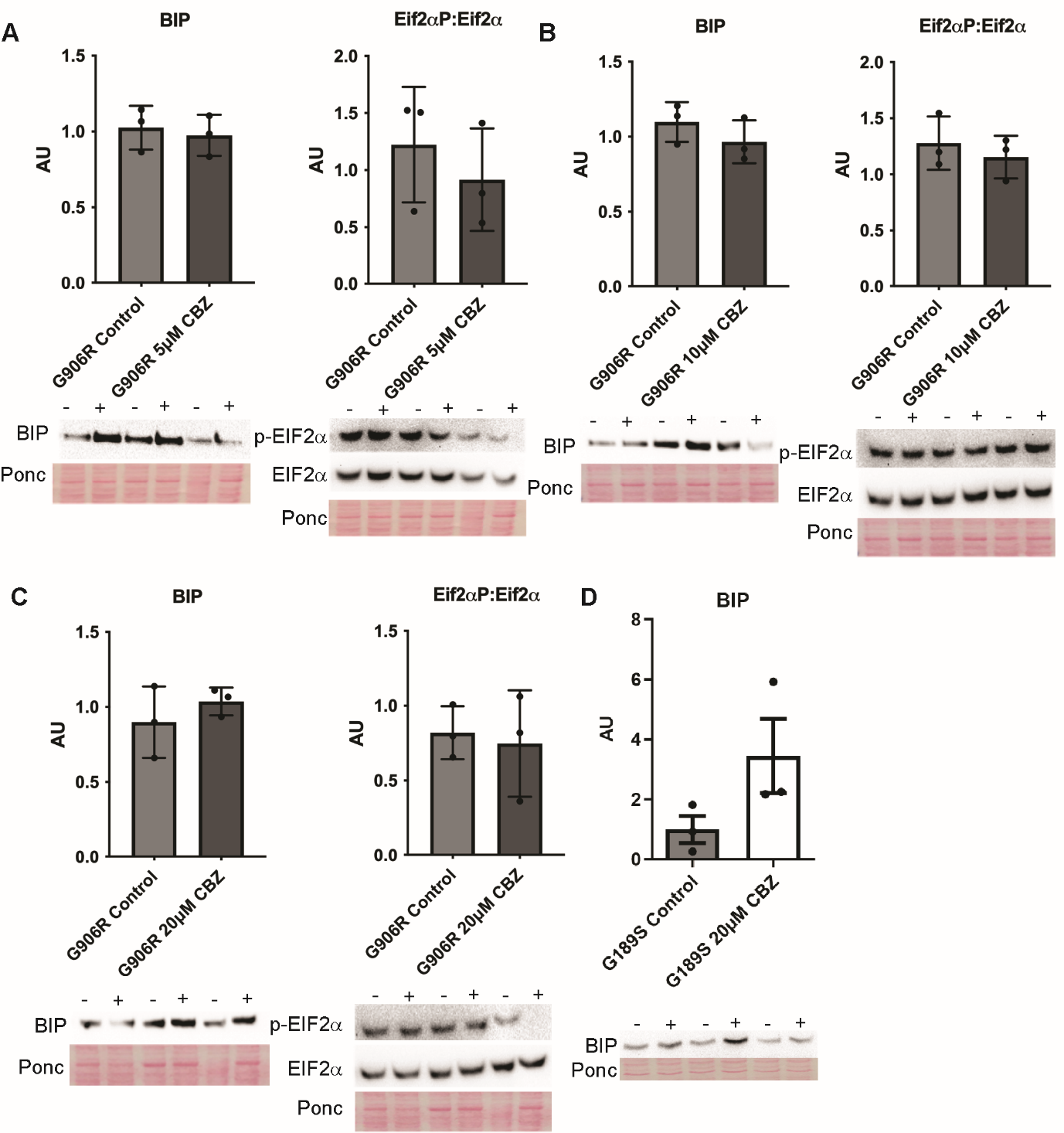

**Supplemental Figure 4. CBZ Treatment does not reduce ER stress levels due to COL3A1 mutations.** Cells were incubated for 24 hours with CBZ at different concentrations and western blotting was performed to assess protein levels of BIP, phosphorylated EIF2α (p-EIF2α) and total EIF2α as markers of ER stress. Ponceau (Ponc) used as protein loading control. (A) Western blot and graph of densitometry analysis of BIP and ratio of p-EIF2α:EIF2α in *COL3A1*^G906R/+^ cells untreated (control) and incubated with 5 µM CBZ. A-D n =3, unpaired t-test (B) Data of incubation with *COL3A1*^G906R/+^ cells untreated (control) and incubated with 10 µM CBZ. (C) Data of incubation with *COL3A1*^G906R/+^ cells untreated (control) and incubated with 20 µM CBZ. (D) BIP protein levels in *COL3A1*^G189S/+^ cells untreated (control) and incubated with 20 µM CBZ.

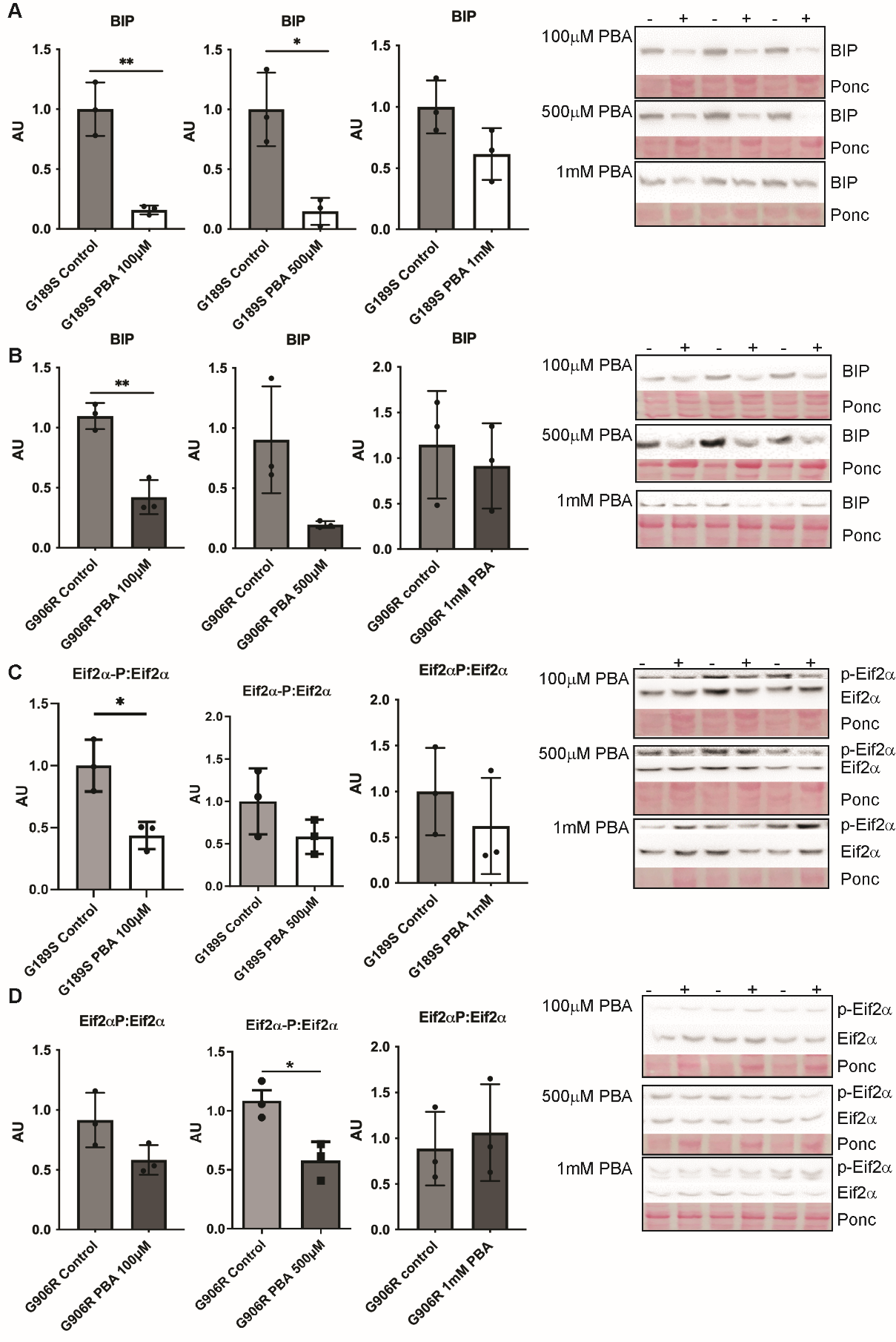

**Supplemental Figure 5. Dosage dependent effects of PBA.** A graphic summary of these data is provided in Figure 4 C-D.(A) BIP protein levels in *COL3A1^G189S^*^/+^ cells treated with PBA for 24 hours compared to untreated cells (control). Western blots are provided on right hand side. (B) BIP protein levels in *COL3A1^G906R^*^/+^ cells treated with PBA for 24 hours compared to untreated cells (control). (C) Ratio of phosphorylated EIF2α (p-EIF2α): total EIF2α in *COL3A1^G189S^*^/+^ cells treated with PBA compared to untreated cells. (D) Ratio of phosphorylated EIF2α (p-EIF2α): total EIF2α in *COL3A1^G906R^*^/+^ cells treated with PBA compared to untreated cells. A-D n=3, unpaired t-test * p<0.05, ** p<0.01

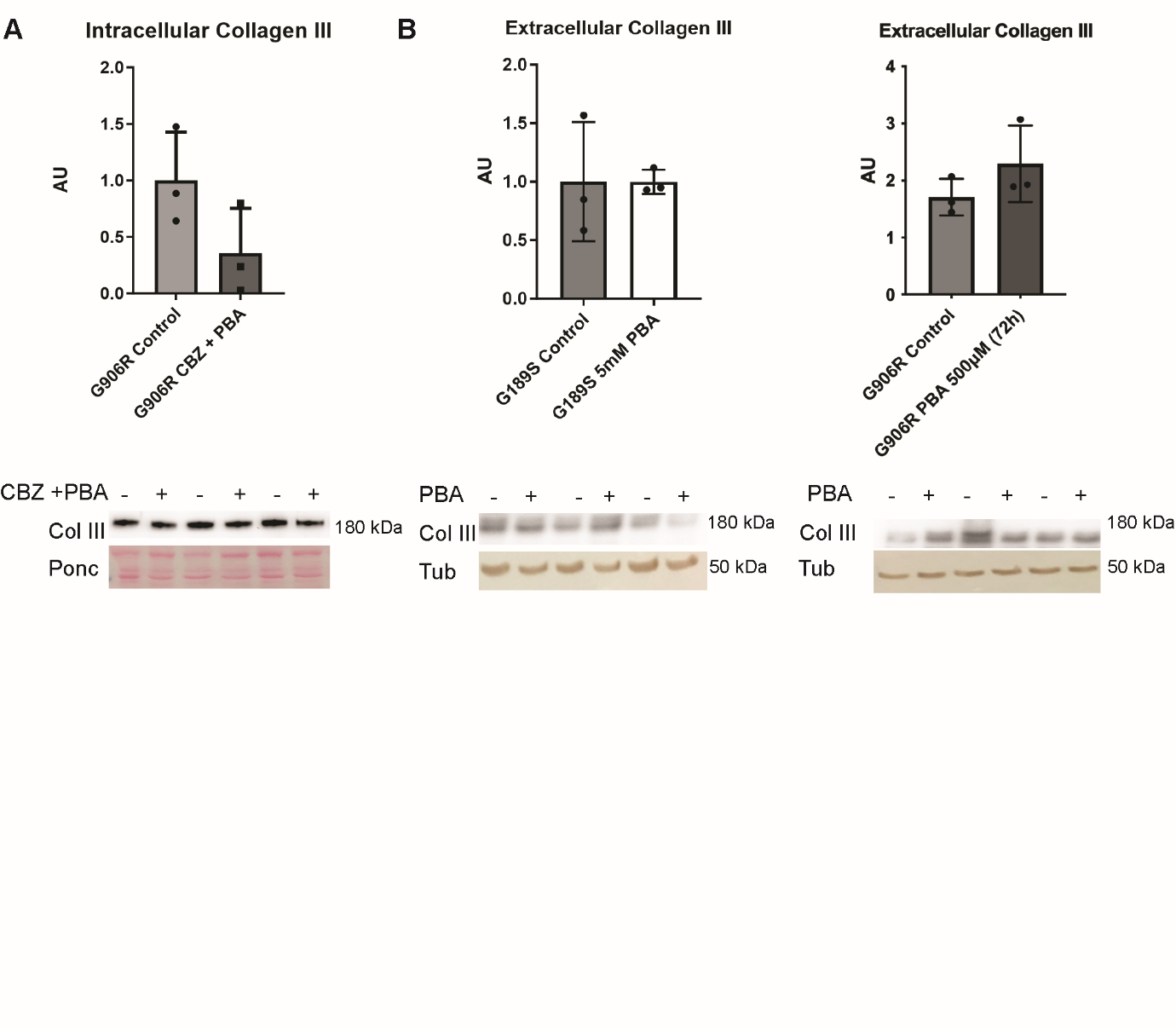

**Supplemental Figure 6. Compound PBA:CBZ treatment and impact of PBA on collagen III secretion.** (A) Western Blot against intracellular collagen III in COL3A1^G906R/+^ cells treated (+) with combinatorial treatment of 500µM PBA and 20µM CBZ for 24 hours. Untreated cells (-, control). No reduction in intracellular collagen III was observed. (n=3, unpaired t-test) (B) Western Blot against collagen III on conditioned media of COL3A1^G189S/+^ cells and COL3A1^G906R/+^ treated (+) with PBA for 24 and 72 hours respectively. Untreated cells (-). Media was changed for the final 24 hours so conditioned media reflected the amount of collagen III secretion over a 24 hour period. No difference in collagen III secretion was observed. (n=3, unpaired t-test).
